# Morphological profiling of environmental chemicals enables efficient and untargeted exploration of combination effects

**DOI:** 10.1101/2022.02.10.479889

**Authors:** Jonne Rietdijk, Tanya Aggarwal, Polina Georgieva, Maris Lapins, Jordi Carreras-Puigvert, Ola Spjuth

## Abstract

Environmental chemicals are commonly studied one at a time, and there is a need to advance our understanding of the effect of exposure to their combinations. Here we apply high-content microscopy imaging of cells stained with multiplexed dyes (Cell Painting) to profile the effects of Cetyltrimethylammonium bromide (CTAB), Bisphenol A (BPA), and Dibutyltin dilaurate (DBTDL) exposure on four human cell lines; both individually and in all combinations. We show that morphological features can be used with multivariate data analysis to discern between exposures from individual compounds, concentrations, and combinations. CTAB and DBTDL induced concentration-dependent morphological changes across the four cell lines, and BPA exacerbated morphological effects when combined with CTAB and DBTDL. Combined exposure to CTAB and BPA induced changes on the ER, Golgi apparatus, nucleoli and cytoplasmic RNA in one of the cell lines. Different responses between cell lines indicate that multiple cell types are needed when assessing combination effects. The rapid and relatively low-cost experiments combined with high information content makes Cell Painting an attractive methodology for future studies of combination effects. All data in the study is made publicly available on Figshare.

**Highlights:**

- Assessment of combination effects of BPA, CTAB and DBTDL on four human cell lines
- Morphological profiling/Cell Painting captures dose and combination dependent effects
- BPA exacerbated morphological effects when combined with CTAB and DBTDL.
- Cell models of diverse origin are needed when profiling environmental chemicals

## 1. Introduction

Much effort is focused on understanding the potential hazard of environmental chemicals. These chemicals can be present in everyday products, such as food and cosmetics, and in our direct environment, such as pesticides, pharmaceuticals and synthetic chemicals^1^. Despite growing scientific evidence for the toxicity of chemical mixtures, little is known about combination effects of environmental chemicals and the possible health risks they pose. Current approaches for risk assessment of chemical mixtures are often based on concentration or response addition models of the individual component toxicities^2^. These methods neglect interactions between chemicals in the environment or at target sites, which can result in overall toxicity being stronger or weaker than predicted^2^. New experimental approaches to test and validate toxicity models to better capture combination effects are greatly needed.

The most widely used methodology to study the cellular response to chemicals in regards to its toxicity or mode of action are viability assays, or alternatively readouts to investigate the activation of specific pathways. Such assays include reactive oxygen species, DNA damage, ER stress, or NFKb activation, among others^1^ and are commonly targeting specific individual biological pathways. It is not uncommon to perform several such assays to get a wider profile of relevant endpoints^3^.

Morphological cell profiling is increasingly being used in diverse scientific areas including toxicology and drug discovery, propelled by the ability to generate rich data at reasonable costs^4,5^. In contrast to traditional single endpoint assays, morphological profiling is untargeted and can be used to study multiple types of cell responses simultaneously. A popular method is Cell Painting^6^ where cells are perturbed with e.g. exposure to a chemical compound or genetic alteration, stained with multiplexed dyes binding to different cell compartments, and imaged using automated microscopy in multiple channels. Image analysis can then be used to segment individual cells, followed by a range of measurements producing morphological profiles on a single-cell level. Contributing to the power and popularity of the method is also that the resulting images and morphological profiles have successfully been analyzed with unsupervised and supervised machine learning methods^7–9^.

Morphological profiling can be used to elucidate toxicity mechanisms and mode of action of a given chemical entity, which in turn allows for the generation of empirical hypotheses^10^. The methodology has been applied to experimental drugs, large chemical libraries, pseudo-natural products, and environmental chemicals across various cell lines^4,7,11–14^. In safety assessment, cell-based assays offer the possibility to provide insights into the cellular responses to chemical exposure. Chemically-induced cellular responses can be cell-specific as well as organelle-specific and are often complex, consisting of initiation and activation of various biochemical processes that lead to a biological response. To this end, morphological profiling and Cell Painting are gaining popularity in screening and profiling applications. However, studies applying morphological profiling to investigate the effects of chemical combinations on cells have so far not been reported.

In this study, we analyze the individual effect as well as combination effects of three environmental chemicals, present in everyday objects and contexts, on four human cell lines using Cell Painting. *Bisphenol A* (*BPA*) is one of the most used plasticizers applied in the polymerization of polycarbonate plastics. BPA has been associated with multiple adverse health effects such as metabolic disorders^15^, cardiovascular diseases^16^, and cancers^17,18^. Due to its ubiquitous presence in consumer products such as food packaging, toys and drinking bottles, this chemical has a high potential to be co-exposed to other chemicals^19^. *Dibutyltin dilaurate* (*DBTDL*) is a widely used industrial chemical, serving as an antifouling coating and is used in pesticides and fungicides. It is a persistent chemical that can be deposited in the liver where it can affect liver functions^20^. *Cetyltrimethylammonium bromide* (*CTAB*) is a cationic compound that is used as an antibacterial and antifungal surfactant, used as a topical antiseptic in cleaning and cosmetic products. This compound has been shown to induce cytotoxicity and inflammatory toxicity *in vivo* and *in vitro*^21^.

The aim of this study is to establish that morphological profiling with Cell Painting combined with multivariate analysis is a suitable methodology that can be used to analyze the effect of combinations of chemicals on cells, demonstrated on three chemical substances that have widespread use.

## 2. Materials and Methods

### Cell culture

U-2 OS cells (ATCC; HTB-96), MCF7 cells (Sigma; 86012803) and A549 cells (ATCC; CCL-185), were cultured in Dulbecco’s Minimum Essential Media (DMEM; Thermo Fisher Scientific; 31885-023) supplemented with 10% (v/v) fetal bovine serum (FBS; Thermo Fisher Scientific; 11550356), 50 U/ml penicillin and 50 μg/ml streptomycin (P/S; Thermo Fisher Scientific; 15140122). Caco-2 cells (ATCC; HTB-37) were cultured in Minimum Essential Medium (MEM; Thermo Fisher Scientific, 31095-029) supplemented with 10% FBS and 1% P/S. The cells were maintained at 37 °C under 5% CO_2_. When the cells reached 80 to 90% confluency, cells were washed with DPBS 1X (Gibco; 14190250), dissociated with TrypLE (Gibco; A1217701) and reseeded in fresh complete medium in T75 flasks (Thermo Scientific; 156499).

### Cell seeding and compound treatments

Using a Biotek Multiflo FX microplate dispenser, U-2 OS (1000 cells/well), A549 (800 cells/well), Caco-2 (1200 cells/well) and MCF7 (1200 cells/well) were seeded in a volume of 24μl in 384 multiwell plates (Costar; Falcon 384-well Optilux Flat Bottom plates, 353962). The plates were kept at room temperature for 20 minutes to aid homogeneous spreading. The plates were incubated overnight for 24 hours at 37 °C at 5% CO_2_ atmosphere to allow for cell attachment. The outer wells were excluded from experimentation to avoid edge effects.

The three selected environmental chemicals in this study included: BPA (Sigma-Aldrich; 239658), DBTDL (Sigma-Aldrich; 291234) and CTAB (Sigma-Aldrich; H9151). The compounds were tested individually, as well as in two and three-component mixtures in three concentrations (1μM, 6.45μM and 12μM). Six phenotypic reference compounds were included which are known to induce a distinctive phenotype in a variety of cells^22^. Specifically, these included: Etoposide (E1383), Fenbendazole (F5396), Metoclopramide (M0763), Berberine Chloride (B3251), Tetrandrine (T2695) and Fluphenazine dihydrochloride (F4765-1G). D-Sorbitol (S1876) was included as negative control and DMSO (D2438) was used as a compound vehicle (all purchased from Sigma Aldrich). All compounds were dissolved in DMSO to 10mM stock solutions, then a 5X source plate was prepared in Dulbecco’s Minimum Essential Media (DMEM) using an automated liquid handler (OT-2; Opentrons, Brooklyn, NY). Compound conditions were distributed over the plates with four technical replicates and two biological replicates per cell line. From the 5X source plate, 6 μl compounds was transferred to the assay plates using a Viaflo 384 electronic pipette, reaching 1 × of the final compound concentration (1μM, 6.45μM and 12μM). The cells were incubated for 24 hours at 37 °C under 5% CO2. In order to optimally distribute the conditions over the plate and reduce the impact of positional effects, we produced plate layouts (**Suppl. Fig. S1**) with PLAID (Plate Layouts using Artificial Intelligence Design, https://github.com/pharmbio/plaid).

### Cell Painting

The Cell Painting experiments closely followed the protocol described by Bray et al. (2016)^6^. In short, upon 24 hours of chemical exposure, the cells were washed three times with 80μl 1X PBS (Thermo Fisher, cat.no 11510546). A washing protocol was set up for the Biotek 405 LS microplate washer leaving 10μl liquid in the wells between and after each washing step to minimize disturbance of the cells. After the first washing step, 35μl MitoTracker (Invitrogen; M22426) in prewarmed FluoroBrite (Thermo Fisher Scientific; 15291866) was applied to the cells (final well-concentration: 900nM) and incubated for 30 minutes at 37°C in a humidified atmosphere under 5% CO2. Then, the MitoTracker solution was removed and the cells were washed three times (80μl, 1x PBS) followed by fixation in 50μl of 4% PFA (Histolab; 02176) for 20 minutes. Plates were washed an additional three times, followed by permeabilization with 50μl of 0.1% Triton X-100 for 20 minutes at room temperature. Triton X-100 was removed followed by three washes with 1X PBS. Then a staining step was performed, including Hoechst 33342 (Invitrogen; H3570), SYTO 14 green (Invitrogen; S7576), Concanavalin A/Alexa Fluor 488 (Invitrogen; C11252), Wheat Germ Agglutinin/Alexa Fluor 555 (Invitrogen; W32464) and Phalloidin/Alexa Fluor 568 (Invitrogen; A12380) in 1X PBS. A total of 20 μl staining mixture was added to each well reaching a final well-concentration of 1μg/ml Hoechst, 15μg/ml Wheat germ agglutinin, 10μl/ml Phalloidin, 4μM SYTO 14 and 80μg/ml Concanavalin A, and was incubated for 20 minutes. Then, the plates were washed a final three times (80μl, 1x PBS), sealed and kept at 4°C prior to imaging. Plates were protected from light as much as possible.

### Image acquisition and processing

Fluorescence microscopy was conducted using a high throughput ImageXpress Micro XLS (Molecular Devices) microscope with a 20X objective with laser-based autofocus. Offsets were determined for each cell line and kept constant throughout the experiment. Six sites per well were captured using 5 fluorescence channels to capture the DNA (Hoechst), mitochondria (MitoTracker), Golgi apparatus and plasma membrane (Wheat Germ Agglutinin), F-actin (Phalloidin), nucleoli and cytoplasmic RNA (SYTO 14), and the endoplasmic reticulum (Concanavalin A/Alexa Fluor 488). Excitation spectra were set to 377/50 nm (Hoechst), 628/40 nm (Mitotracker), 562/40 nm (Phalloidin and Wheat germ agglutinin), 531/40 nm (SYTO 14) and 482/35 nm (Concanavalin A). Emission filters were set to detect signals between 447–60 nm (Hoechst), 692/40 nm (MitoTracker), 624/40 nm (Wheat Germ Agglutinin and Phalloidin), 593/40 nm (SYTO 14) and 536/35 (Concanavalin A). Images were processed and analyzed with the open-source image analysis software CellProfiler version 4.0.6 23 and CellPose generalist algorithm for cellular segmentation^24^.

Image analysis was divided into three steps: 1) quality control, 2) illumination correction and 3) segmentation and feature extraction. The quality control (QC) pipeline was run on the raw images to detect images with staining and/or imaging artifacts. Various quality measures were calculated on the raw images to represent a wide variety of artifacts. Images deviating more than five standard deviations from the median for FocusScore, MaxIntensity, MeanIntensity, PercentMaximal, PowerLogLogSlope and StdIntensity were flagged, inspected and removed if necessary. Blurred and oversaturated images were detected by calculating the PercentMaximal score and PowerLogLogSlope. Images with PercentMaximal values higher than 0.25, or PowerLogLogSlope values lower than −2.3 were removed from the dataset. The remaining bright artifacts in the DAPI channel were identified using the Identify Primary Object module in CellProfiler. The artifacts and surrounding pixels were masked to avoid interference with segmentation. Intensity measures, object size and cell counts were visualized as plate heatmaps to detect plate and batch effects. Images with a cell count below 20 were removed from further analysis. To correct for uneven illumination, a polynomial illumination correction function was calculated for each plate and each image channel. Each image was then divided by the respective illumination correction image to correct for uneven illumination.

A feature extraction pipeline was built to segment three cell compartments (nuclei, cytoplasm, cells) and extract morphological features. Nuclei segmentation was performed on the Hoechst staining, by applying a gaussian blur followed by Otsu thresholding to segment the outlines of each nucleus. For U-2 OS, A549 and MCF7, the cell objects were segmented using watershed segmentation using minimum cross-entropy on the cytoplasmic RNA stain, using the nuclei as a seed. Size and thresholds for nuclei and cytoplasm segmentation were adjusted for each cell line. For Caco-2 cells, composites of the Hoechst and Phalloidin+WGA staining were used for cellular segmentation using the CellPose cyto segmentation model. The resulting cell masks were loaded in CellProfiler and cell objects were related to the corresponding nuclei located within the area of each cell mask. The cytoplasm compartment was defined as the cell object subtracted from the nuclei.

Phenotypic characteristics were measured for the three cell compartments, using the AreaShape, Correlation, Intensity, RadialDistribution, Granularity, Location and Neighbors, modules as provided by CellProfiler. A total of 2330 features were extracted from each cell, which were exported into CSV format for downstream analysis. Image analysis pipelines (quality control, illumination correction and feature extraction) are made publicly available in Figshare.

### Multivariate analysis

Downstream analysis was performed using Python 3. First, the median was computed for all features on an image level. Varying and outlier features (SD < 0.001 and SD > 10000), as well as features with missing values were removed. The image-based feature values were mean centered and normalized to unit variance using the fit.transform method of StandardScaler of scikit-learn module (version 0.22.1). Perturbation-level profiles were computed by aggregating to replicate-level profiles by computing their mean. The high dimensional data was visualized in lower dimensional space using principal component analysis (PCA) on the centered and normalized features and Uniform Manifold Approximation and Projection (UMAP) using sklearn (version 0.22.1). UMAP visualization was done using 30 or 40 neighbors. Partial least-squares discriminant analysis (PLS-DA) was used to model compound-induced morphological profiles. Q2 and R2 values were calculated for the PLS-DA models, which were estimated by three-fold cross-validation repeated for 10 times. Induction scores were computed using z-score normalization relative to the DMSO controls per plate: x.zscore = (x – median(x[subset]))/mad(x[subset]). The induction score was used as a measure for bioactivity, as described earlier^25^. Specifically, it represents the fraction of features (in %) that was significantly changed (median absolute deviation (MAD) upon compound treatment of at least +/- 1.96 of the median determined for the DMSO controls). For visualization of the affected features in radar plots, the absolute mean of the z-score normalized features was computed, grouped by CellProfiler module (Intensity, Correlation, Granularity, Location, RadialDistribution), and stain(s) (Hoechst, Concanavalin A, SYTO 14, Mitotracker, WGA and Phalloidin). Areashape and Neighbors features were grouped per cell compartment.

### Data availability

All 33,600 images of the exposed four cell lines (U-2 OS, MCF7, A549, and Caco-2) together with CellProfiler image analysis pipelines, extracted morphological features, and metadata (204 GB in total) are deposited at Figshare: https://doi.org/10.17044/scilifelab.19063028.

## 3. Results

### 4.1 Evaluation of environmental toxicants by morphological profiling

In order to investigate the morphological effects the environmental toxicants CTAB, BPA and DBTDL (**Fig. 1A**) exert on cells, we used the Cell Painting assay^6^. In short, cell lines originally derived from different organ origin, represented by MCF7 (breast cancer), A549 (lung cancer), Caco-2 (colon cancer) and U-2 OS (osteosarcoma), were seeded on multiwell plates 24h prior to exposure to combinations of the environmental toxicants for another 24h. Subsequently, the Cell Painting protocol was performed, followed by high-content imaging, image analysis using Cell Profiler and CellPose^23,24^, and data analysis of the morphological features (**Fig. 1B**). To assess the reproducibility and sensitivity of the assay, we first analyzed the morphological changes induced by DMSO controls and six phenotypic reference compounds. Principal component analysis (PCA) of the morphological profiles in the DMSO condition separated each cell line in the morphological space, and are highly reproducible between biological replicates (**Fig. 1D**). Exposure to the phenotypic reference compounds induced dose-dependent changes that showed substantial similarity between cell lines (**Suppl. Fig. S2**). Next, we assessed the effect of exposing the four cell lines to the environmental toxicants CTAB, BPA and DBTDL. The cell lines responded differently to each of the treatments, which is depicted by the morphological changes observed upon the staining of the different cell compartments included in the Cell Painting assay (**Fig. 1C**). In particular, distinct morphological changes could be observed upon CTAB exposure on A549 and Caco-2 cells, characterized by intra-cytoplasmic vesicles, as well as increased intensity of mitochondrial staining upon DBTDL exposure for all cell lines (**Fig. 1C**). We generated morphological profiles of MCF7, A549, Caco-2 and U-2 OS cells exposed to vehicle (DMSO), CTAB, BPA, and DBTDL, at 1, 6 and 12 μM, for 24h (**Suppl. Figs. S3 and S4**).

**Figure 1:**
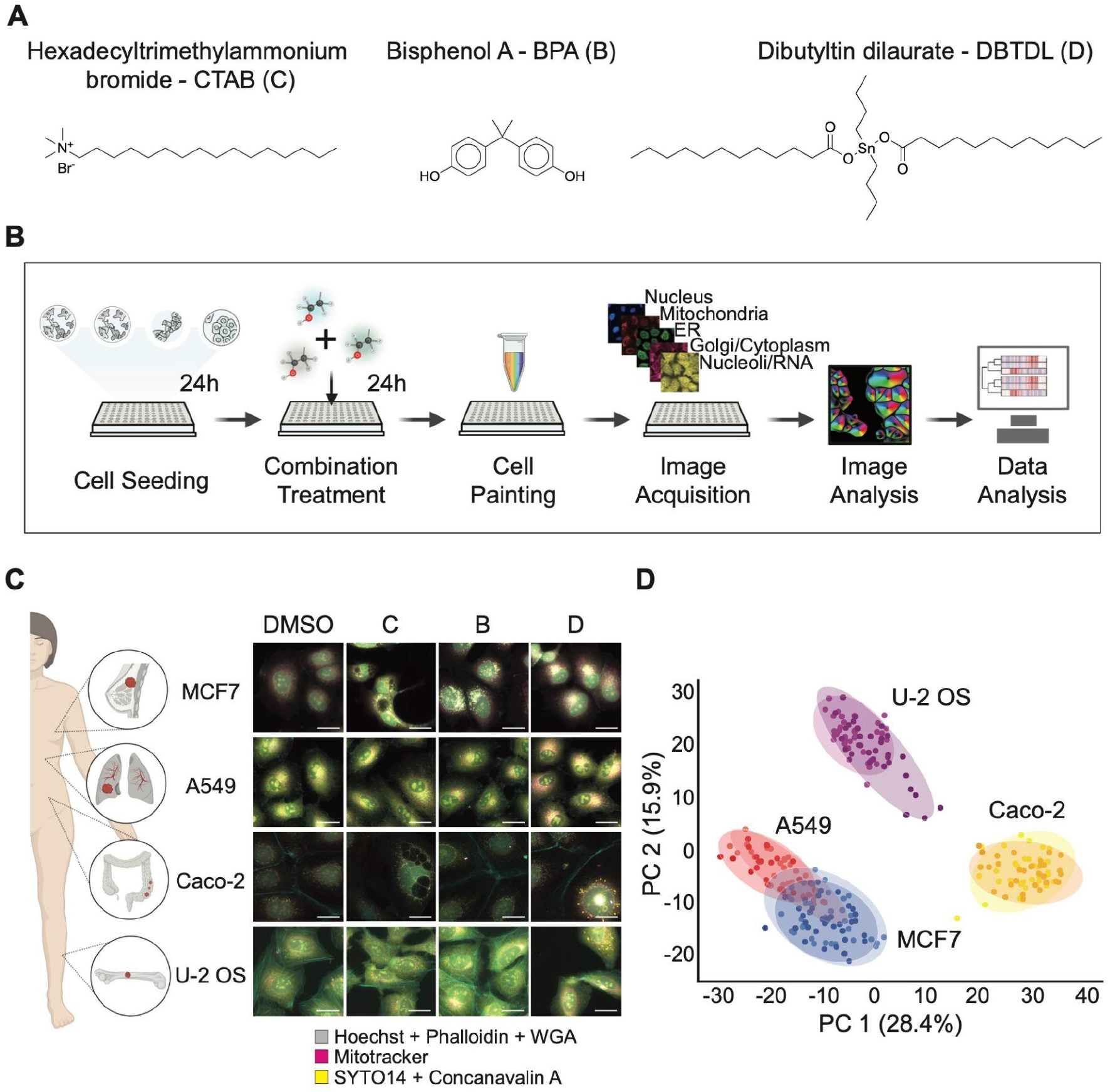
Morphological profiling of chemical combinations in cells of four different origins. **A**. Chemical structure of the three environmental chemicals used in this study. **B**. Overview of the morphological profiling approach. Cells are seeded in multiwell plates 24h prior to exposure to the environmental chemicals, either at single, double or triple combinations. After 24h of exposure, automated Cell Painting, high content imaging, image analysis and data analysis is performed. **C**. Representative images of the four cell lines used in this study, MCF7 breast cancer cells, A549 lung cancer cells, Caco-2 colon cancer cells and U-2 OS osteosarcoma cells, with Hoechst (Nuclei), Phalloidin (F-actin) and Wheat Germ Agglutinin (Golgi apparatus) in gray, Mitotracker (Mitochondria) in red and SYTO14 (Nucleoli and cytoplasmic RNA) and Concanavalin A (ER) in yellow. Cells were exposed to vehicle (DMSO), C, B, or D, at 6μM for 24h. Scale bar is 25μm. **D**. Principal Component Analysis (PCA) of the morphological profiles of MCF7, A549, Caco-2 and U-2 OS cells in DMSO for the two biological replicates (indicated by color shade). Illustrations were partially created with BioRender.com. C: CTAB; B: BPA; D: DBTDL.

### 4.2 CTAB, BPA and DBTDL induce cell-specific morphological signatures

We used cell count to assess the effect of the chemical exposures at 1, 6 and 12μM (**Fig. 2A**). The highest concentration of CTAB and DBTDL seem to induce toxicity in all cell lines, however DBTDL particularly affected U-2 OS cells. In order to quantitatively assess morphological changes, we calculated an induction score (IS) based on the z-score of the number of features altered by the exposure to each of the compounds, in respect to the vehicle DMSO (**Fig. 2B**). The IS clearly showed a dose-dependent increase in the alteration of the morphological features for DBTDL and CTAB, but to a much lesser extent for BPA.

**Figure 2:**
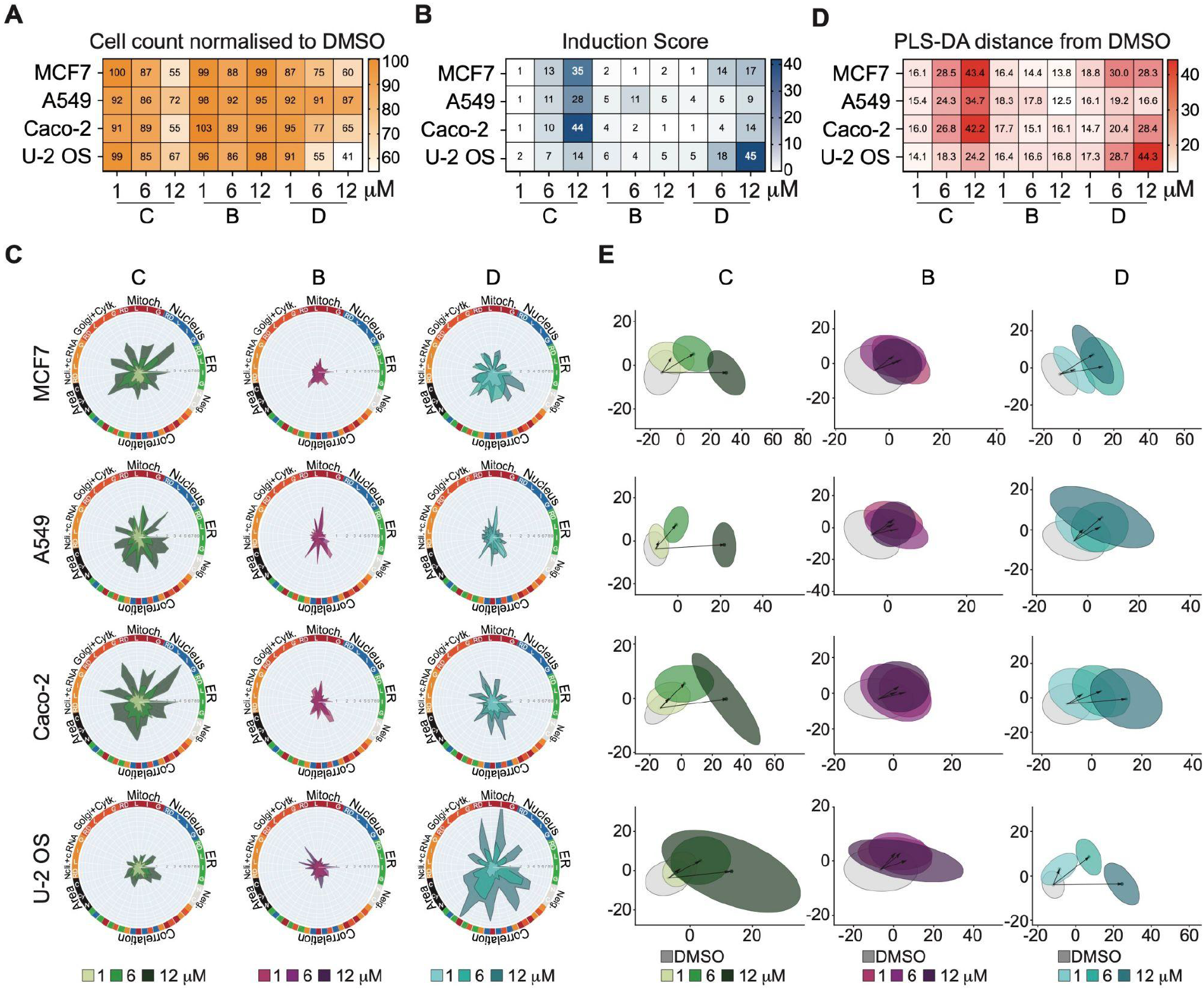
Morphological profiling captures cell line-specific and chemical-specific phenotypic signatures. **A**. Cell count heat map of MCF7, A549, Caco-2 and U-2 OS cells exposed to C, B, and D at 1, 6 or 12μM for 24h, normalized to the vehicle (DMSO). Lower number of cells is represented by a lighter color. **B**. Induction Score heat map of cells treated as in **A**, representing the percentage of affected features relative to DMSO. Darker color corresponds to a higher number of features affected by the exposure. **C**. Radar plots for each of the conditions as in **A**, representing the affected morphological features. The features were grouped by Cell Profiler module, i.e. Intensity (I), Correlation (C), Granularity (G), Location (L) and RadialDistribution (RD); as well as by stains, i.e. Nucleus (Hoechst), ER (Concanavalin A), Nucleoli and cytoplasmic RNA (SYTO14), Golgi apparatus and F-actin cytoskeleton (WGA and Phalloidin) and Mitochondria (Mitotracker). Area-shape related features were grouped by cell compartment, i.e. Cell (C), Cytoplasm (Cy) and Nucleus (N). Neighboring related features were grouped by Cell (C) and Nucleus (N) and Neighbour features were grouped per cell compartment, i.e. Correlation features among the different stains were represented by their corresponding color code. **D**. Calculated PLS-DA distances between the median of the samples corresponding to each condition as in **A** and DMSO. Darker color corresponds to cellular morphologies increasingly deviating from DMSO controls. **E**. PLS-DA scatter plots of the samples corresponding to each condition as in **A**; ellipses correspond to 90% confidence interval; arrows indicate the distance from medians of DMSO towards the medians of the indicated treatments. C: CTAB; B: BPA; D: DBTDL.

Next, in order to explore what cellular compartments were altered upon exposure to the three environmental chemicals, we constructed radar plots by calculating the z-score of the extracted morphological features, grouped by cellular compartment (nucleus, endoplasmic reticulum (ER), nucleoli and cytoplasmic RNA, F-actin cytoskeleton and Golgi apparatus, and mitochondria), as well as correlation, neighboring and area features, for each cell line, chemical and dose (**Fig. 2C**). The radar plots for CTAB exposure indicated a clear dose-response effect on all cell lines. CTAB induced a prominent effect on all cellular compartments in all cell lines, with the exception of U-2 OS, which displayed a lower response. Features related to granularity and intensity, as well as area of both nuclei and cells, were particularly affected by CTAB exposure. BPA, seemingly induced a lower response in all cell lines, except for A549 in which the intensity of the mitochondria was sharply increased at exposure, and U-2 OS in which the intensities of nuclei, cytoplasmic RNA, Golgi apparatus and F-actin cytoskeleton were increased. Exposure to DBTDL resulted in an increased granularity of all cellular compartments in MCF7 and Caco-2 cells, but to a lesser extent in the case of A549 cells. In contrast, DBTDL induced a pronounced increase in the intensity of the mitochondria staining, as well as changing in correlation features between mitochondria and ER, nucleus and the Golgi apparatus and F-actin cytoskeleton in U-2 OS cells.

We used partial least-squares discriminant analysis (PLS-DA) on the extracted morphological features to create models for predicting the effects that each compound elicited on the cells. Similarly to the Induction Scores, the PLS-DA distance from DMSO (**Fig. 2D**), indicated a dose-response for CTAB and DBTDL, which was not the case for BPA. The PLS-DA models predicted that CTAB and DBTDL induced distinct and reproducible morphological changes, indicated by the clustering of the different chemical doses, distanced from the DMSO control, as well as the high coefficient prediction Q^2^, especially for concentrations 6μM and 12μM (**Fig. 2E and Supplementary Table S1**). On the other hand, the low coefficient prediction Q^2^ for BPA for most concentrations (**Fig. 2E and Supplementary Table S1**), indicated that the morphological changes induced by this chemical did not differ substantially from the DMSO control, as it is also seen by the close proximity of the clusters of the three BPA concentrations to DMSO (**Fig. 2E**).

### 4.3 Cell Painting enables detection of combination effects

We next investigated the effect of the combination of CTAB, DBTDL and BPA. We exposed MCF7, A549, Caco-2 and U-2 OS cells to either double or triple combinations of the chemicals, at 1, 6 or 12μM. We used cell count to determine the toxicity resulting from combined exposure to the chemicals (**Fig. 3A**). Only at the highest concentration, either double or triple exposure of the compounds seemingly induced some degree of toxicity. With the exception of the combination of CTAB and DBTDL, BPA and DBTDL, as well as the triple CTAB+BPA+DBTDL combination, which induced toxicity in MCF7, Caco-2, and particularly on U-2 OS cells, at 6μM (**Fig. 3A**).

**Figure 3:**
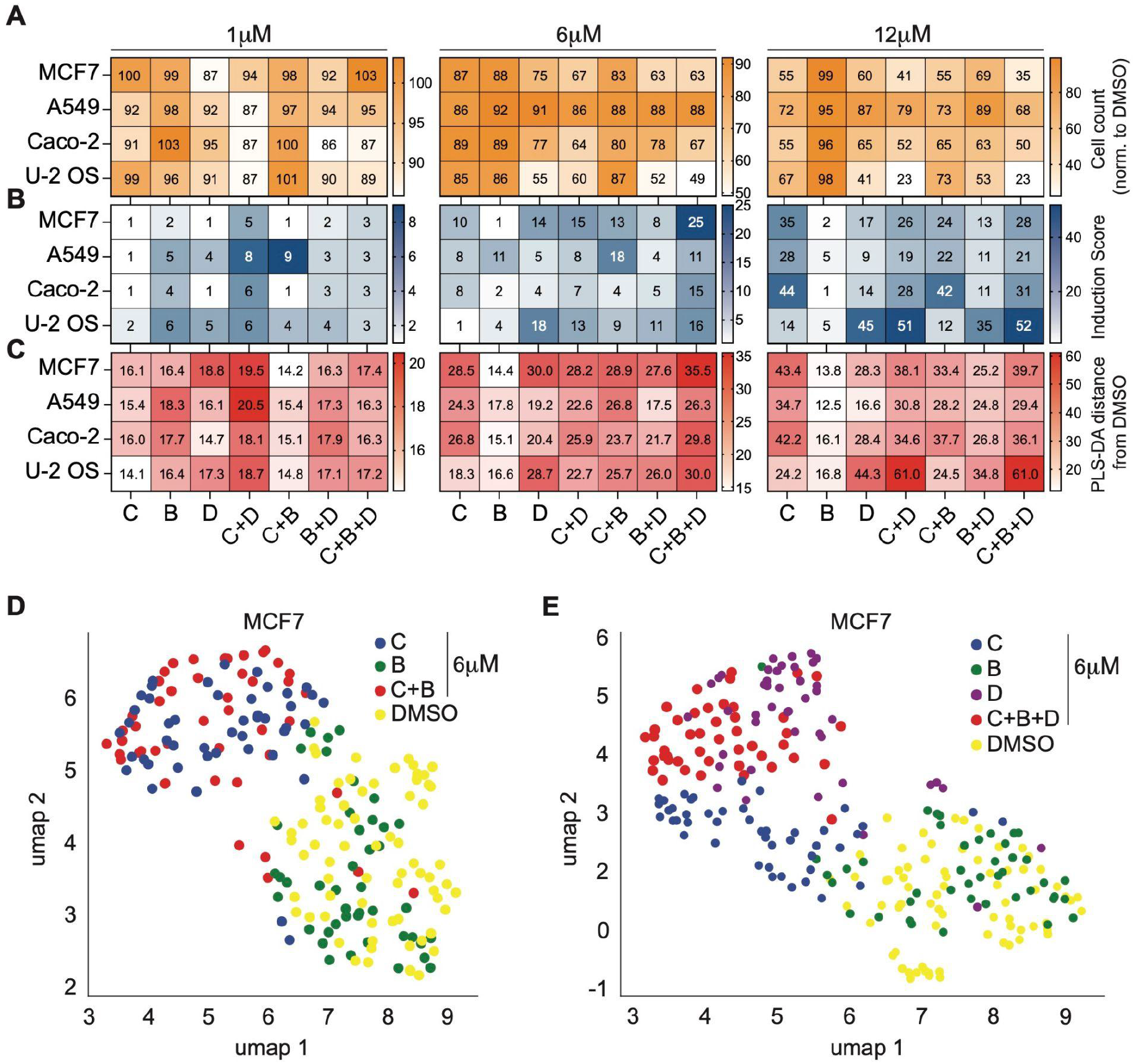
Morphological profiling captures phenotypic signatures of chemical combinations. **A**. Cell count heat maps of MCF7, A549, Caco-2 and U-2 OS cells exposed to C, B, and D at 1, 6 or 12μM for 24h, normalized to the vehicle (DMSO), as well as the indicated combinations (C+D, C+B, B+D and C+B+D). Lower number of cells is represented by a lighter color. **B**. Induction Score heat map of cells treated as in **A**, representing the percentage of affected features relative to DMSO. Darker color corresponds to a higher number of features affected by the exposure **C**. Calculated PLS-DA distances between the median of the samples corresponding to each condition as in **A** and DMSO. Darker color corresponds to cellular morphologies increasingly deviating from DMSO controls. Of note, the values for the single exposure conditions are already shown in **Fig. 2**, but displayed here too for clarity. **D**. Uniform manifold approximation and projection (UMAP) on the median of the morphological profiles of MCF7 cells exposed to DMSO, C, B or combined exposure to C+B, for 24h at 6μM. **E**. UMAP on the image median of the morphological profiles of MCF7 cells exposed to DMSO, C, B, D or combined exposure to C+B+D, for 24h at 6μM. C: CTAB; B: BPA; D: DBTDL.

We calculated the Induction Scores (IS) for both double and triple combinations of chemicals (**Fig. 3B**). The IS indicated that combining DBTDL with either CTAB or BPA at 6μM, resulted in a reduction of the score compared to the exposure of the chemicals alone, pointing at a potential buffering effect (**Fig. 3B**). The combination of CTAB and BPA at 6μM and the combination of DBTDL and CTAB at 1μM, however, resulted in an increased IS compared to the single chemical exposures in U-2 OS, A549 and MCF7 cells, indicating a potential additive or synergistic effect. PLS-DA was then used on the extracted morphological features from the double and triple exposures. The PLS-DA distances from DMSO reflected a similar effect as identified by the Induction Score calculation, here we observed a shorter distance from DMSO when exposing the cells to DBTDL in combination to CTAB compared to the individual exposures (**Fig. 3C**).

To further explore the cell morphologies induced by combined exposure, the features were dimensionally reduced using non-linear uniform manifold approximation and projection (UMAP). (**Fig. 3D, 3E**). Morphological profiles induced by combined exposure to CTAB and BPA were dominated by the effects from CTAB in MCF7 cells (**Fig. 3D**). Triple exposure to compounds CTAB, BPA and DBTDL, induced morphological changes that could be attributed to the cumulative changes from both CTAB and DBTDL (**Fig 3E**).

The radar plots representation of the morphological features confirmed the results of the Induction Score calculation for the combination treatment with CTAB and DBTDL in U-2 OS and MCF7 cells, in which the double exposure to these chemicals resulted in a reduction of the morphological alterations compared to the single exposures at 6μM (**Fig. 4A**). In contrast, exposure to CTAB and BPA resulted in either exacerbation of the morphological changes induced by the single exposure to the chemicals, or in the alternation of morphological features not altered by the single exposures, such as alteration of the ER intensity in U-2 OS cells. This double combination induced perturbation of the ER, Golgi apparatus, Nucleoli and cytoplasmic RNA in A549 cells (**Fig. 4A**), which is depicted by a decrease in intensity of the stainings in the corresponding images (**Fig. 4B**).

**Figure 4:**
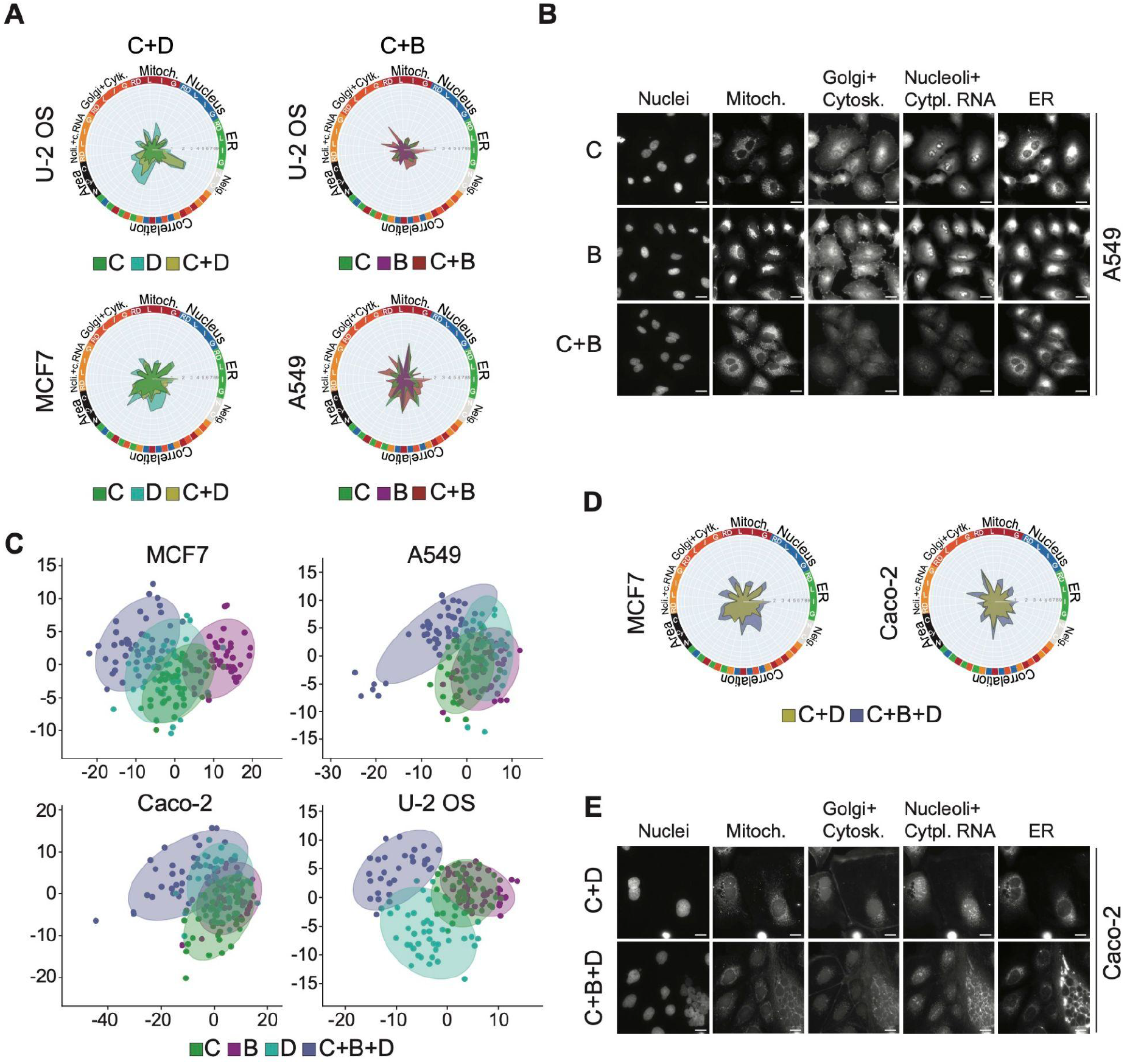
Chemical combinations induce distinct responses. **A**. Radar plots for single or combined exposure to C, D, B, C+D or C+B in U-2 OS and MCF7 cells, at 6μM for 24h, representing the affected morphological features. **B**. Representative images of A549 cells exposed to C, BPA or combined exposure to C+B at 6μM for 24h. Scale bar is 20μm. **C**. PLS-DA scatter plots of MCF7, A549, Caco-2 and U-2 OS cells exposed to C, B, D or C+B+D, at 6μM for 24h. Ellipses indicate 90% Confidence Interval. **D**. Radar plots for combined exposure to C+D or C+B+D in MCF7 and Caco-2 cells. At 6μM for 24h. **E**. Representative images of Caco-2 cells exposed to the combination of C+D or C+B+D, at 6μM for 24h. Scale bar is 20μm. C: CTAB; B: BPA; D: DBTDL.

PLS-DA analysis comparing single to triple chemical exposures resulted in a separation between the conditions in all cell lines, in particular in the case of A549 and U-2 OS cells, where the triple exposure clustered separately from all the single exposures (**Fig. 4C**).

An increase in global perturbation levels of the morphological features was observed by the addition of BPA, compared to the double combination of CTAB and DBTDL. Addition of BPA to DBTDL and CTAB, resulted in larger PLS-DA distances (**Fig. 3B**) and higher induction scores (**Fig. 3C**) across all four cell lines at 6μM and 12μM exposures. The radar plots display increased perturbation levels in MCF7 and Caco-2 cells upon addition of BPA (**Fig. 4D** and **E**).

## 4. Discussion

The continued increase in the number and load of chemicals in our environment highlights the need for methods to systematically assess risks associated with exposure to combinations. Here, we employed morphological profiling as a new experimental approach to study combination effects of chemicals. We interrogated chemical-induced cellular morphologies of three environmental chemicals: CTAB, DBTDL and BPA exposed individually as well as in double and triple combinations, at three different doses, using four human cell lines.

Using the Cell Painting assay with single-cell measures of cell morphologies and multivariate data analysis, we were able to identify distinct changes upon exposure to both single chemicals and combinations of chemicals. We could effectively discern effects from individual doses for chemicals CTAB and DBTDL, as well as distinguish between single and combined exposure with CTAB, DBTDL and BPA. BPA did not result in significant morphological changes but showed additive or synergistic effects when co-exposed to CTAB and DBTDL.

Exposure to compounds CTAB and DBTDL followed a clear concentration-dependent morphological response across all four tested cell lines. In contrast, exposure to compound BPA resulted in a more subtle morphological response, which followed a non-monotonic dose-dependent response. This result could be explained by opposing effects induced by compound BPA binding to multiple receptors, receptor desensitization, negative feedback loops, or dose-dependent metabolism modulation, which are commonly reported for endocrine-disrupting chemicals^26^.

Previous research has shown that DBTDL can induce cell cycle alterations, DNA damage, and increased calcium influx, and at higher concentrations can lead to nuclei deformation, mitochondrial swelling, and apoptotic-like changes^27,28^. In concordance with these findings, we observed that high concentrations of DBTDL heavily affected the mitochondria in all cell lines, as well as the nuclei intensity and size of nuclei and cell compartments in U-2 OS and MCF7 cells, which might be an indication for apoptosis.

CTAB induced strong morphological changes in all cell organelles and was bioactive in at least three out of the four cell lines. This finding is in line with previous observations where CTAB present in gold nanorods has been shown to induce cell apoptosis and autophagy by activating reactive oxygen species (ROS) and leading to mitochondrial dysfunction^29^. One proposed mechanism of toxicity of compound CTAB, like other surfactants, is the formation of micelles, also known as nanobubbles, in aqueous solutions. These micelles can disrupt cell membranes leading to Ca^2+^ influx and subsequent cell death and can induce inflammatory responses when exposed to human blood cells ^30^.

Interestingly, even though compound BPA did not elicit a strong morphological change when exposed alone, an increase in perturbation level could be observed when combined with CTAB or when combined with both compounds DBTDL and CTAB together. Previously, co-exposure of BPA with other endocrine-disrupting chemicals, such as phthalates, have been reported to lead to synergistic and additive interactions, such as increased DNA damage *in vitro*, and significantly changed lipid profiles and glucose levels *in vivo*^31,32^. The increased bioactivity could also be explained by interactions between BPA and CTAB molecules; BPA can stabilize CTAB molecules in the micelle and at the air/water interface which might in turn increase the micelle-mediated toxicity by CTAB ^33^.

Another important finding in this study is the observed cell-line specific effects upon treatment. We used breast cancer MCF7 cells, lung cancer A549 cells, colon cancer Caco-2 cells, and osteosarcoma U-2 OS cells, all of which differ on their genetic, metabolic or proteomic landscapes^34–36^. Thus, the diverse responses to the environmental chemicals by the different cell lines could be explained by the presence or absence of the specific chemicals’ molecular targets. This finding emphasized the need to include biologically diverse cell lines for toxicity studies.

One surprising result in this study was the dose-dependent dampening or exacerbation of morphological effects for combined exposure of CTAB and DBTDL. Commonly used methods, such as additive and response-addition models only assume additive effects upon combined exposure, which further highlights the need for experimental tools to validate toxicity models. However, to develop a full picture of the combination effects between the chemicals, additional studies will be needed exploring a wider range of concentrations and using concentration-response matrices.

Morphological profiling has the ability to capture subtle morphological changes upon a given perturbation, before it reaches toxic levels. This indicates that we can detect cell stress upon exposure to contaminants with cell morphology profiling before it can be observed in e.g. cell viability assays. Further, compared to widely used receptor-based assays, morphological profiling captures a much wider set of changes at molecular and cellular levels integrated in a single assay. Indeed, analyses of Cell Painting profiles have earlier been shown to be predictive for multiple cell health outcomes such as proliferation, apoptosis, reactive oxygen species (ROS), DNA damage, cytotoxicity, and cell cycle phase ^9,37^.

The untargeted and unbiased approach makes Cell Painting an attractive alternative to receptor based assays to test higher order environmental chemical mixtures. However, one possible shortcoming is assay sensitivity to detect changes at low concentrations. In this study we observed only subtle changes at the 1μM concentration, whereas environmentally relevant concentrations of exposure of environmental contaminants are commonly much lower^1^. Possible explanations could be that assay variability and batch effects mask subtle morphological effects, that the multivariate analysis approach does not capture certain relevant changes, or that the biological activity does not manifest in a morphological change. In addition, the use of primary cell lines, or non-immortalised cell lines, might improve the detection of the subtle morphological changes that environmental toxicants may induce. Integration of additional profiling techniques, such as proteomics or transcriptomics, may shine more light on cellular responses.

Humans are potentially exposed to more than thirty thousand chemicals from a variety of sources, screening even a fraction of all possible combinations would be impossible^19^. Risk assessment of chemical mixtures is a multifaceted task, which should integrate experimental toxicity data, mechanism of action and in silico modeling^38^. Major bottlenecks for mixture models nowadays are that experimental toxicity data is scattered and insufficient to address complex mixtures. In this study we show that morphological profiling offers a reproducible and standardized approach, integrating multiple endpoints and cell lines, which could be used to systematically assess combination effects and has the potential to advance our understanding of health risks of chemical mixtures.

## Abbreviations

(CTAB): Cetyltrimethylammonium bromide
(DBTDL): Dibutyltin dilaurate
(BPA): Bisphenol A
(IS): Induction Score

